# The expression of equine keratins K42 and K124 is restricted to the hoof epidermal lamellae of *Equus caballus*

**DOI:** 10.1101/678102

**Authors:** Caitlin Armstrong, Lynne Cassimeris, Claire Da Silva Santos, Yagmur Micoogullari, Bettina Wagner, Susanna Babasyan, Samantha Brooks, Hannah Galantino-Homer

## Abstract

The equine hoof inner epithelium is folded into primary and secondary epidermal lamellae which increase the dermo-epidermal junction surface area of the hoof and can be affected by laminitis, a common disease of equids. Two keratin proteins (K), K42 and K124, are the most abundant keratins in the hoof lamellar tissue of *Equus caballus*. We hypothesize that these keratins are lamellar tissue-specific and could serve as differentiation- and disease-specific markers. Our objective was to characterize the expression of K42 and K124 in equine stratified epithelia and to generate monoclonal antibodies against K42 and K124. By RT-PCR analysis, keratin gene (*KRT*) *KRT42* and *KRT124* expression was present in lamellar tissue, but not cornea, haired skin, or hoof coronet. In situ hybridization studies showed that *KRT124* localized to the suprabasal and, to a lesser extent, basal cells of the lamellae, was absent from haired skin and hoof coronet, and abruptly transitions from *KRT124*-negative coronet to *KRT124*-positive proximal lamellae. A monoclonal antibody generated against full-length recombinant equine K42 detected a lamellar keratin of the appropriate size, but also cross-reacted with other epidermal keratins. Three monoclonal antibodies generated against N- and C-terminal K124 peptides detected a band of the appropriate size in lamellar tissue and did not cross-react with proteins from haired skin, corneal limbus, hoof coronet, tongue, glabrous skin, oral mucosa, or chestnut on immunoblots. K124 localized to lamellar cells by indirect immunofluorescence. This is the first study to demonstrate the localization and expression of a hoof lamellar-specific keratin, K124, and to validate anti-K124 monoclonal antibodies.

## 1. Introduction

The skin and its appendages are made of stratified epithelia composed of keratinocytes, defined by expression of keratin intermediate filament proteins (abbreviated K for proteins and *KRT* for genes) [1]. Keratin filaments resist stretching (strain) and provide tensile strength to epithelia and skin appendages. Tissue- and differentiation-specific variation in specific keratin isoform content determines the physical and mechanical properties of diverse epithelial tissues and of their keratinocyte building blocks [2;3]. Understanding how keratins function to provide mechanical stability is crucial to our understanding of human diseases associated with keratin mutations and abnormal keratin expression [4-8]. Here we describe unique keratins of the equine (*Equus caballus*) epidermal lamellae, a highly specialized tissue that withstands extreme force and provides a model for understanding how keratins provide mechanical strength to flexible tissues.

Each single-toed foot of a 500 kg horse (*E. caballus*) must withstand peak ground reaction forces of 2-5,000 N while protecting the underlying limb from trauma [9;10]. The equine adaptation to single-toed unguligrade locomotion requires the integration of the musculoskeletal system with a cornified hoof capsule and the strong, but flexible, suspension of the distal phalanx from the inner surface of the hoof capsule [11-13]. As shown in Fig 1, the inner epithelium of the equine hoof capsule, which is homologous to the nail bed [14], is folded into primary and secondary epidermal lamellae (PELs and SELs, respectively), thus increasing the surface area of epidermal-dermal attachment and, with the dermal connective tissue to which it adheres, forming the suspensory apparatus of the distal phalanx (SADP) [12]. Structural failure of the SADP results in laminitis, a common and crippling disease of equids and other ungulates [15]. In spite of the importance of the hoof lamellae for equine locomotion and disease, few aspects of hoof lamellar biology, including keratin isoform composition, have been well characterized.

**Fig 1:**
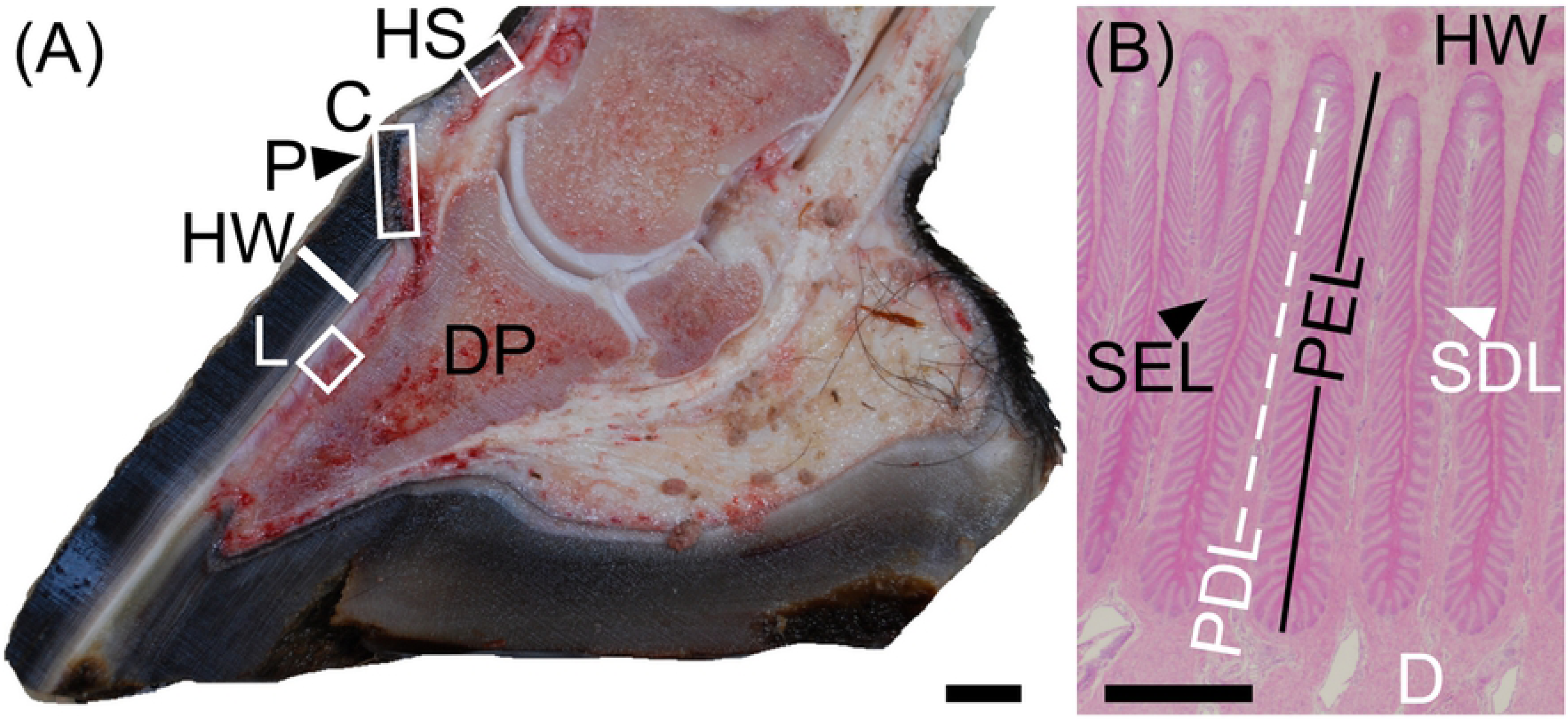
Macroscopic anatomy of the equine (*E. caballus*) foot and microscopic anatomy of the hoof lamellae. (**A**) Equine foot, midsagittal section, showing locations of samples retrieved for this study: HS: haired skin and the following hoof capsule regions: C: coronet (proximal stratum medium layer and nail matrix homolog), P: periople (stratum externum layer and cuticle homolog), HW: hoof wall (stratum medium layer and nail plate homolog), L: lamellar tissue, including epidermal lamellae (stratum internum layer and nail bed homolog), dermal lamellae, and dermal corium up to the surface of the distal phalanx (DP). (**B**) Transverse section of the lamellar region (H&E stain). HW, at the top of the image, is contiguous with the approximately 500 cornifying primary epidermal lamellae (PELs) of each hoof capsule. Each PEL has 100-150 secondary epidermal lamellae (SELs; black arrowhead). The PELs and SELs interdigitate with primary dermal lamellae (PDLs) and secondary dermal lamellae (SDLs) which are in turn continuous with the dermal corium (D), transferring the weight of the horse from the DP to the HW. Scale bars: A: 1 cm; B: 500 μm

Equine keratin isoform expression and localization has relied entirely on commercial antibodies, many of which cross-react with multiple keratin isoforms [16;17]. Similar to other stratified epithelia, hoof lamellae express K14 in the basal cell layer, and also express unique keratin isoforms that contribute to the health and disease of this tissue [18;19]. By proteomics, we discovered two novel equine keratins, K42 and K124.[18] These keratins are the most abundant cytoskeletal proteins in equine hoof lamellae, accounting for over fifty percent of the total keratin content of this tissue [18].

*KRT42* and *KRT124* exist only as pseudogenes in humans, *KRT42P* and *KRT90P*, respectively (the latter was formerly named *KRT124P* when equine *KRT124* was named [1;18;20]). Murine *Krt42* (formerly *K17n* or *Ka22*) mRNA is expressed in the nail unit [21]. A putative *KRT124* ortholog, *Krt90* (formerly *Kb15*), is translated from cDNA libraries in mice and rats [22] and was identified from the draft genomic sequence of the opossum [23]. *KRT42* and *KRT124* were recently identified and mapped to the canine and equine genomes, but their patterns of expression have not been described beyond their identification from RNA-seq data derived from skin biopsies from three dogs and one horse [20]. The lack of isoform-specific antibodies has impeded the detailed investigation of equine hoof capsule or lamellar tissue-specific keratins [19]. The objectives of this study were to characterize the pattern of expression of K42 and K124 in equine stratified epithelia of the hoof and skin and to determine if the most abundant keratins of the hoof lamellae are specific differentiation markers of this highly specialized epithelium. We report here that K42 and K124 expression is restricted to equine hoof lamellae and we have characterized monoclonal antibodies against K124.

## 2. Methods

### 2.1 Ethics statement

The protocols, titled ‘Pathophysiology of Equine Laminitis (By-products only)’ and ‘Equine Laminitis Tissue Bank,’ under which the archived equine tissue samples used for this study were collected, were approved by the University of Pennsylvania Institutional Animal Care and Use Committee (protocol #801950 and #804262, respectively). Euthanasia of the horses was carried out in accordance with the recommendations in the Guide for the Care and Use of Agricultural Animals in Research in Teaching Federation for Animal Science Societies) and the AVMA Guidelines for the Euthanasia of Animals (American Veterinary Medical Association) by overdose with pentobarbital sodium and phenytoin sodium. Mouse immunization, euthanasia by cervical dislocation, and monoclonal antibody production were carried out in accordance with the recommendation in the Guide for the Care and Use of Laboratory Animals of the National Institute of Health. The protocol titled ‘Mouse Monoclonal Antibody Production’ was approved by the Institutional Animal Care and Use Committee at Cornell University (protocol #2007-0079).

### 2.2 Subjects and tissue retrieval

All *E. caballus* subjects are part of a laminitis tissue repository, were euthanized for medical reasons unrelated to this study, as previously described [24], and had no clinical history, macroscopic or microscopic evidence of hoof, dermatological, or corneal diseases in the tissues used [25]. Age, breed, sex, and tissues used from each subject are listed in S1 Table. Anatomical locations of tissues dissected from the foot are illustrated in Fig 1A. Haired skin, coronet (coronary region of the hoof, homologous to the nail matrix [14], including epidermal and supporting dermal tissue at the proximal edge of the hoof capsule, from which the hoof wall (nail plate) grows), and lamellar tissues (the innermost layer of the hoof capsule, homologous to the nail bed [14], including PELs and SELs, corresponding primary and secondary dermal lamellae and adjacent dermal corium) were collected immediately after euthanasia, as described elsewhere [24;26;27]. All other tissues were dissected by scalpel immediately post mortem. Tissue samples were immediately either 1) snap frozen in liquid nitrogen and stored in liquid nitrogen until processed for protein or RNA extraction, 2) formalin-fixed/paraffin-embedded (FFPE) until sectioned for in situ hybridization studies, or 3) paraformaldehyde-fixed/sucrose-dehydrated, embedded, frozen, and stored at −80°C until sectioned for indirect immunofluorescence, as previously described [18;28].

### 2.3 Oligonucleotide primers, RNA extraction, and qualitative PCR

Oligonucleotide primers for equine keratin isoforms *KRT10A, KRT10B, KRT14, KRT42*, and *KRT124* were designed using Primer3 (http://bioinfo.ut.ee/primer3-0.4.0/)[29] and are listed in Table 1. All oligonucleotides were synthesized by Integrated DNA Technologies, Inc. (Coralville, IA, USA). Alternate forward primers were designed to amplify the two *KRT10* genes, *KRT10A* and *KRT10B*, since the equine *KRT10* gene has undergone duplication [20]. For *KRT124*, one set of primers was designed to amplify the 3’ exon (*KRT124 3’*), which shows no homology to known keratins, and a second primer set was designed to amplify a region of the predicted transcript that shows some homology to several keratins (*KRT124 Mid*). For all primers, RT-PCR product sequence was confirmed by Sanger sequencing for at least one band from each positive tissue type (data not shown).

**Table 1:**
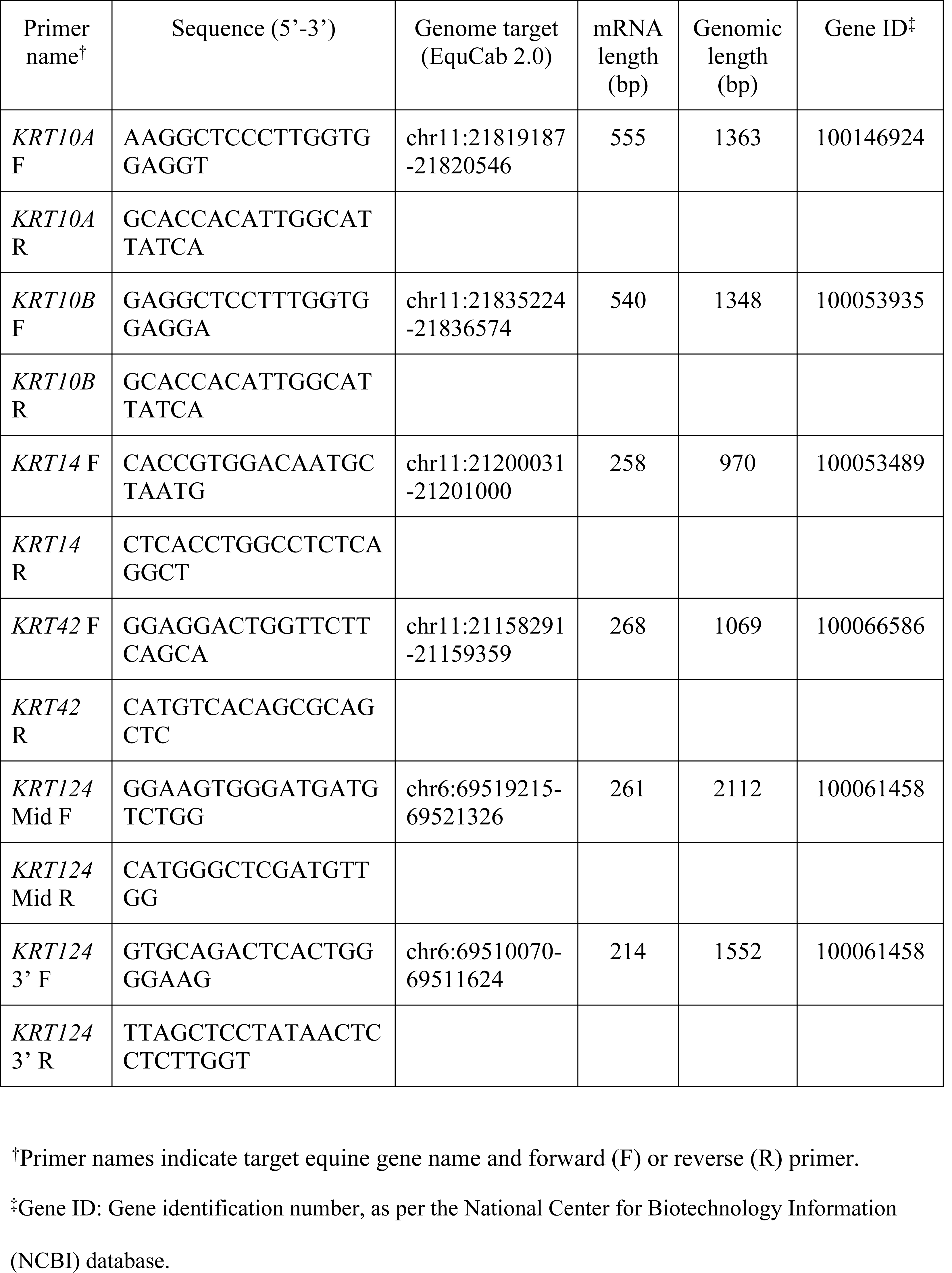
PCR Primer sequences.

#### RNA extraction

Archived snap-frozen tissue was pulverized using a liquid nitrogen-chilled, ELIMINase-treated (Decon Labs Inc, King of Prussia, PA, USA) stainless steel mortar and pestle (Bio-pulverizer™, BioSpec Products, Bartlesville, OK, USA) prior to total RNA extraction. Next, total RNA was extracted using the RNeasy Fibrous Tissue Mini Kit (Qiagen, Valencia, CA, USA), by a modified version of the manufacturer’s instructions, as follows, to allow for complete disruption and homogenization of the highly fibrous lamellar tissue. Pulverized tissue samples were mixed with 300 µl Buffer RLT, 590 µl RNase-free water, and 10 µl proteinase K by gentle vortexing, and incubated at 55°C for 10 min. This mixture was then homogenized by repeated (5-10x) aspiration through an 18 g hypodermic needle and syringe and subjected to centrifugation at 10,000 × g for 10 min. The resulting supernatant was then removed and mixed with 0.5 volume of 100% ethanol, added to the provided RNeasy Mini column in 700 µl increments, and subjected to centrifugation 10,000 × g. RNA retained on the RNeasy column was retrieved by adding 350 µl Buffer RW1 and subjecting it to centrifugation for 15 s at 10,000 × g. DNase treatment and subsequent buffer RW1 and RPE steps were then performed according to the manufacturer’s instructions, including the optional final centrifuge step. Final RNA elution was done once in a total volume of 50 µl of RNase-free water. Total RNA was quantified using a Nanodrop 2000 spectrophotometer (Thermo Fisher Scientific, Waltham, MA, USA). Quality was confirmed by checking A260/A280 ratios and ensuring they were 2.0±0.1 for all samples tested.

#### Qualitative RT-PCR

Reverse transcriptase-polymerase chain reaction (RT-PCR) was carried out on an Eppendorf Mastercycler thermocycler (Eppendorf, Hamburg, Germany) in a single step using the Qiagen OneStep RT-PCR Kit (Qiagen), according to the manufacturer’s instructions, in a total reaction volume of 25 µl using 100 ng of total RNA. The RT reaction was done at 50°C for 30 min. The PCR cycling conditions which immediately followed were as follows: 95°C activation step (15 min, 1×) followed by 30 cycles of 3-step PCR (94°C denature for 30 sec, 50°C anneal for 30 sec, 72°C extension for 1 min). The *KRT42* product was produced with the same overall method above but with the following modifications: 66°C anneal; 40 PCR cycles; 2 min extension per cycle with a final 10 min extension (also at 72°C) following completion of all 40 PCR cycles. RT-PCR products were subjected to agarose gel electrophoresis, visualized with SYBR Safe™DNA stain (Thermo Fisher Scientific, Waltham, MA, USA) and imaged on a UV transilluminator with a digital camera and ethidium bromide filter (Canon G10, Tokyo, Japan).

### 2.4 In situ hybridization

Digoxigenin (DIG)-labeled riboprobes that included unique DNA sequences encoding *KRT14* and *KRT124* were produced by gene synthesis. Unique DNA sequences encoding *KRT14* and *KRT124* were produced by gene synthesis. For *KRT14*, a 297 bp region at the 3’ end of the mRNA sequence (NM_001346198; bp 1371-1667) was synthesized. To facilitate cloning, 5’ Not1 and 3’ Kpn1 restriction sites were included in the synthesized DNA fragment. For *KRT124*, a specific sequence of 684 bp was synthesized that included the 3’ end of the coding sequence and a portion of the 3’ UTR (XM_001504397.3; bp 1623-2307). 5’ Not1 and 3’ Xho1 restriction sites were included in the DNA synthesis. The synthesized DNAs were cloned into pBluescript SK (+) vectors at the multi-cloning site, which includes T7 and T3 RNA polymerase promoter sequences. Plasmid construction was verified by Sanger sequencing (data not shown). All gene synthesis and cloning were performed by Genscript (genscript.com). Digoxigenin (DIG)-labeled riboprobes were synthesized from T7 (for antisense probe synthesis) and T3 (for sense probe synthesis) promoters using MEGAscript Transcription kits (Ambion, Thermo Fisher Scientific) according to the manufacturer’s protocol.

In situ hybridization was performed as described, using standard methods [30]. Briefly, FFPE tissue sections were deparaffinized for 2×10 min in xylene, followed by rehydration in a graded ethanol series (100%, 75%, 50% and 25%, 3 min each) and digested for 5 min with proteinase K (10μg/mL; Ambion). Tissue sections were allowed to hybridize overnight in a humid chamber at 65°C with 1 ng/µL of sense (negative probe) or antisense (positive probe) DIG-labeled riboprobes in hybridization buffer containing 50% formamide. After washes in saline-sodium citrate buffer, the sections were incubated with alkaline phosphatase-conjugate anti-DIG Fab fragments (#11093274910, 1:5000, SigmaAldrich, St. Louis, MO, USA) in a humid chamber overnight at 4°C. After washing in PTB (Phosphate Buffered Saline + 0.2% Triton x-100 + 0.1% BSA), labeled probe was visualized using NBT/BCIP substrate (Roche Diagnostics, Indianapolis, IN, USA) resulting in a blue/purple precipitate. PTw buffer (Phosphate Buffered Saline + 0.1% Tween) was used to stop the reaction. Sections were mounted in 80% glycerol/PTw. Images were collected on a Nikon Nti microscope using a Nikon DS /Ti2 color camera and Nikon Elements software (Nikon Instruments, Inc., Melville, NY, USA). Typically images were collected at 10X (brightfield) or at 20X (DIC optics).

### 2.5 Monoclonal antibodies

#### K42 mAb

The entire coding sequence of *KRT42* (gene ID: 100066586) was amplified using gene-specific primers and the total RNA, RT-PCR, and agarose gel electrophoresis methods described in section 2.2. These primer sequences were as follows:

ATGGCTGCCACCACCACCAC (forward primer) and GCGATGGCTGCCCCTTGA (reverse primer). The corresponding genomic location is chr11 + 21153825-21160809 (EquCab2.0). Following excision of the band from the agarose gel, K42 was expressed as a fusion protein with equine IL-4 as previously described [31]. In brief, the 1416-bp product was sub-cloned into the mammalian expression vector (pcDNA3.1 (−)/Myc-His, version B, Invitrogen,Carlsbad, CA, USA) containing equine IL-4 (eIL-4) [31], sequenced for correctness, and used to transiently transfect ExpiCHO-S cells, as per manufacturer’s instructions (Thermo Fisher Scientific). The serum-free cell culture supernatant was harvested after 6 days of incubation and rIL-4/K42 fusion protein was purified, using a HiTrap NHS-Activated HP affinity column coupled with aIL-4 monoclonal antibodies and an ÄKTA Fast Protein Liquid Chromatography (FPLC) instrument (GE Healthcare, Piscataway, NJ, USA). Immunizations, subsequent cell fusion, and mAb screening and selection were performed as previously described [31-33]. Briefly, one BALB/c mouse was immunized with 2 μg purified rIL-4/K42 fusion protein initially followed by 4 injections every 2-3 weeks of 1 μg protein with an adjuvant (Adjuvant MM, Gerbu, Heidelberg, Germany). Three booster injections of 1 μg of rIL-4/K42 without adjuvant were performed prior to euthanasia. Monoclonal antibodies (mAbs) were generated by fusion of splenic B cells from the immunized mouse and murine myeloma cells, as previously described [32].

#### K124

K124 mAbs were produced by Genscript (genscript.com, Piscataway, NJ, USA) by immunizing Balb/c mice with synthetic peptides targeting N-terminal and C-terminal regions of equine K124 (gene ID: 100061458) conjugated to keyhole limpet hemocyanin immunogen followed by splenic lymphocyte fusion with myeloma type SP2/0 cells. Three anti-K124 mAbs were evaluated in our laboratory, and are designated here as K124A (clone 9H8G1, murine isotype IgG2a), targeting the 14 amino acid peptide, (SVSQGGKSFGGGFG) from positions 36-49 of the N-terminal region, and K124C (clone 4G6E9, murine isotype IgG1) and K124D (clone 4G7A3, murine isotype IgG2b), targeting (RIISKTSTKRSYRS), the last 14 amino acids (508-521) of the C-terminal region. Unpurified hybridoma supernatant was used for all K124 mAb experiments.

### 2.6 Immunoblot analysis

Total protein extraction and concentration determination were performed as previously described [18] from the following snap frozen tissues: hoof lamellar, haired skin, and hoof coronet, corneal limbus, chestnut (an epidermal callus on the medial foreleg proximal to the carpus), tongue, oral mucosa, and preputial unhaired (glabrous) skin. SDS-PAGE and immunoblotting were performed as previously described [18], with 2-8 μg total protein loaded per lane and the following dilutions of mouse mAbs: anti-K14 (1:500; clone LL002, Abcam Inc., Cambridge, UK), anti-K42 (1:500), or anti-K124, clones K124A, K124C, or K124D (1:10) followed by secondary goat-anti-mouse-horse radish peroxidase (HRP; 1:5,000, Jackson ImmunoResearch, Inc, West Grove, PA, USA) and chemiluminescence detection by incubation for 1 min with 79 μM p-coumaric acid and 500 μM luminol mixed 1:1 with 3.6 × 10^−3^% hydrogen peroxide, both in 100 mM TRIS-HCl, pH 8.5 (all reagents: Sigma-Aldrich) followed by exposure to x-ray film (Hyperfilm™ECL, GE Healthcare) for 1-3 min and x-ray film development. K124 and K42 immunoblots were reprobed with anti-keratin K14 and mouse anti-β-actin-HRP mAb (K124 blots; 1:15,000, clone AC-15; Sigma-Aldrich) or anti-β-actin-HRP alone (K42 blots) without stripping to demonstrate equal protein load. Following immunoblotting, proteins were visualized by staining with Amido Black staining solution (Sigma-Aldrich) according to manufacturer’s directions.

### 2.7 Immunofluorescence

Indirect immunofluorescence using fluorescein-conjugated wheat germ agglutinin (F-WGA, Vector Laboratories, Burlingame, CA, USA) as a counterstain on paraformaldehyde-fixed/sucrose-dehydrated/optimal cutting temperature compound (OCT)-embedded frozen tissue sections was performed as previously described, with the following modifications [28]. Antigens were unmasked through 20 min in Antigen Unmasking Solution (Vector Laboratories Inc.) in a 100°C steam bath, followed by cooling to RT on ice. Sections were next submerged for 15 min in 0.1M glycine and four min on ice, followed by 20 min in Background Buster (Innovex Biosciences Inc., Richmond, CA, USA) at RT. Unpurified mouse anti-K124 mAb, clone K124C (1:10 dilution) served as primary antibody and goat anti-mouse Alexa Fluor™594 antibody (1:500 dilution; Invitrogen, Thermo Fisher Scientific) as the secondary antibody. Primary and secondary antibody incubations were for 1h at 23°C in a humidified chamber. All antibodies were diluted in PBS containing 2% normal goat serum (Jackson ImmunoResearch). All wash steps following incubation with the primary antibody were done in PBS/0.05% Tween-20. Sections were mounted and imaged by confocal microscopy as previously described [28]. Primary antibody was omitted to determine background staining.

## 3. Results

### 3.1 *KRT42* and *KRT124* mRNA is detected in hoof lamellae, but not haired skin, cornea, or hoof coronet

As shown in Fig 2, RT-PCR was performed to determine the qualitative tissue expression of the major keratin isoforms that we had previously detected as proteins in equine hoof lamellar tissue, K42 and K124, in the cornea, haired skin, hoof coronet and lamellar tissues. All RT-PCR products display molecular weights predicted to correspond to mRNA rather than genomic DNA (Fig 2, Table 1). The basal cell keratin, *KRT14*, was used as a positive control since it is expressed in all stratified epithelial tissues [34]. *KRT14* is expressed in cornea, haired skin, and lamellae, *KRT10A* and *KRT10B* expression is restricted to haired skin and of the two, *KRT10B* is more readily detected by this method. *KRT42* and *KRT124* mRNA is only detected in lamellar tissue and was not amplified from mRNA isolated from cornea, haired skin, or hoof coronet tissues.

**Fig 2:**
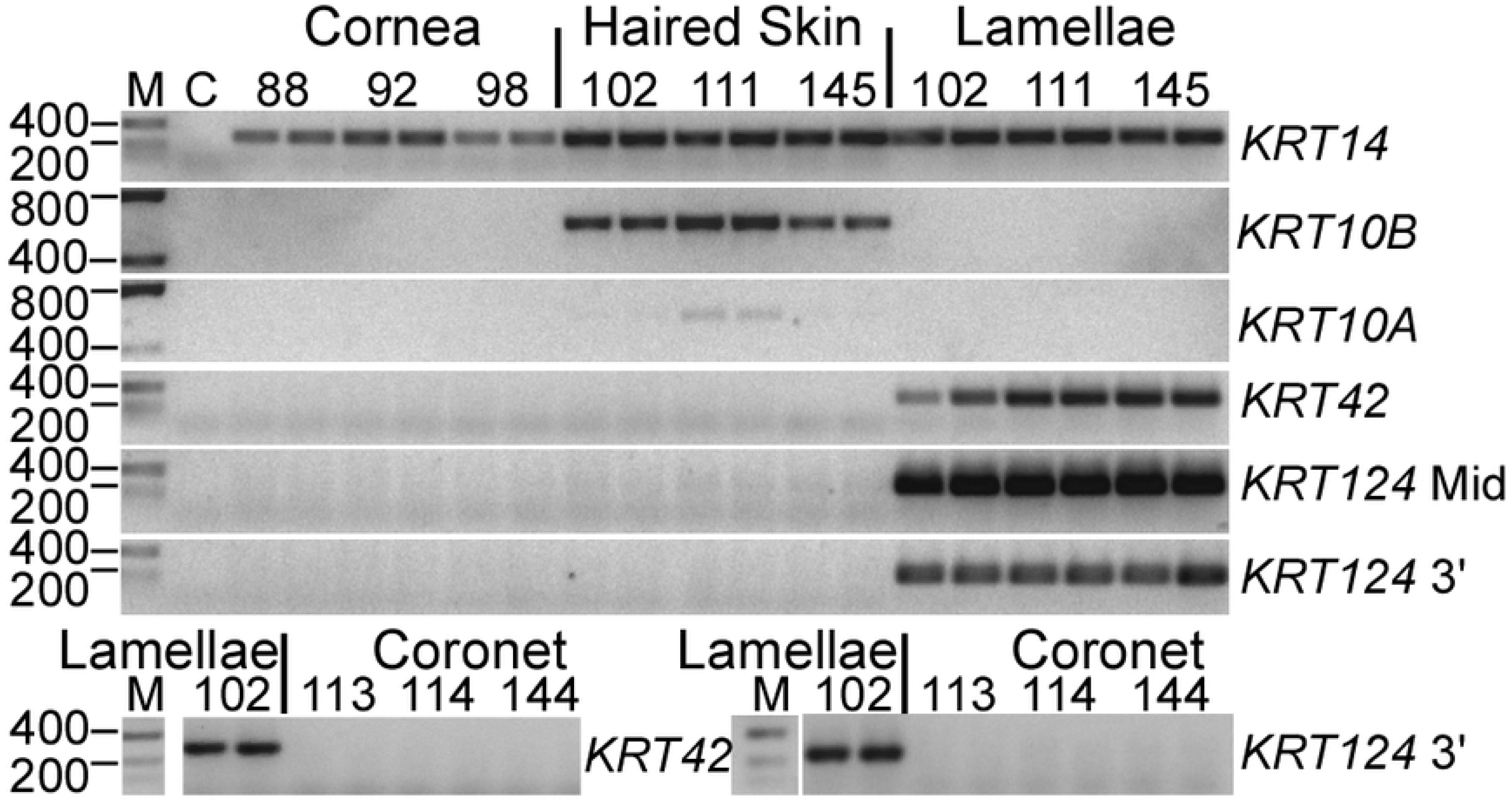
*KRT42* and *KRT124* are expressed in hoof lamellae, but not cornea, haired skin, and coronet. Representative RT-PCR products from equine cornea, haired skin, hoof lamellae, and hoof coronet, using primers for *KRT14, KRT10A, KRT10B, KRT42*, and *KRT124*, as indicated to the right of gels, and separated by agarose gel electrophoresis, produces amplicons of the expected base pair sizes. RT-PCR products from duplicate experiments were run using RNA extracts from three different horses (identified by number above pairs of lanes) per tissue. DNA ladder (M), negative control without template RNA (C), and tissues identified above gels. Duplicate *KRT10* genes present in separate loci that were individually amplified using specific primers (*K10A* and *K10B*). Two sets of primers were used to amplify two different regions of *KRT124* (*KRT124* Mid and *KRT124* 3’). Image inverted for ease of viewing.

### 3.2 *KRT124* mRNA localizes to the hoof secondary epidermal lamellae and is absent from hoof coronet and haired skin

As shown in Fig 3-4, we employed in situ hybridization (ISH) to more precisely localize *KRT124* expression. *KRT124* was detected by ISH in all regions of the epidermal lamellae except for the central keratinized axis of the primary epidermal lamellae (Fig 3A). *KRT124* expression was not detected in the coronet or in haired skin (Fig 3B). Staining was also negative in lamellar tissue using a *KRT124* sense probe (Fig 3A, lower panels). An abrupt transition from *KRT124*-negative coronary epidermal tissue to *KRT124*-positive lamellar epidermal tissue is apparent at the junction between coronet and the first proximal lamella (Fig 3B).

**Fig 3:**
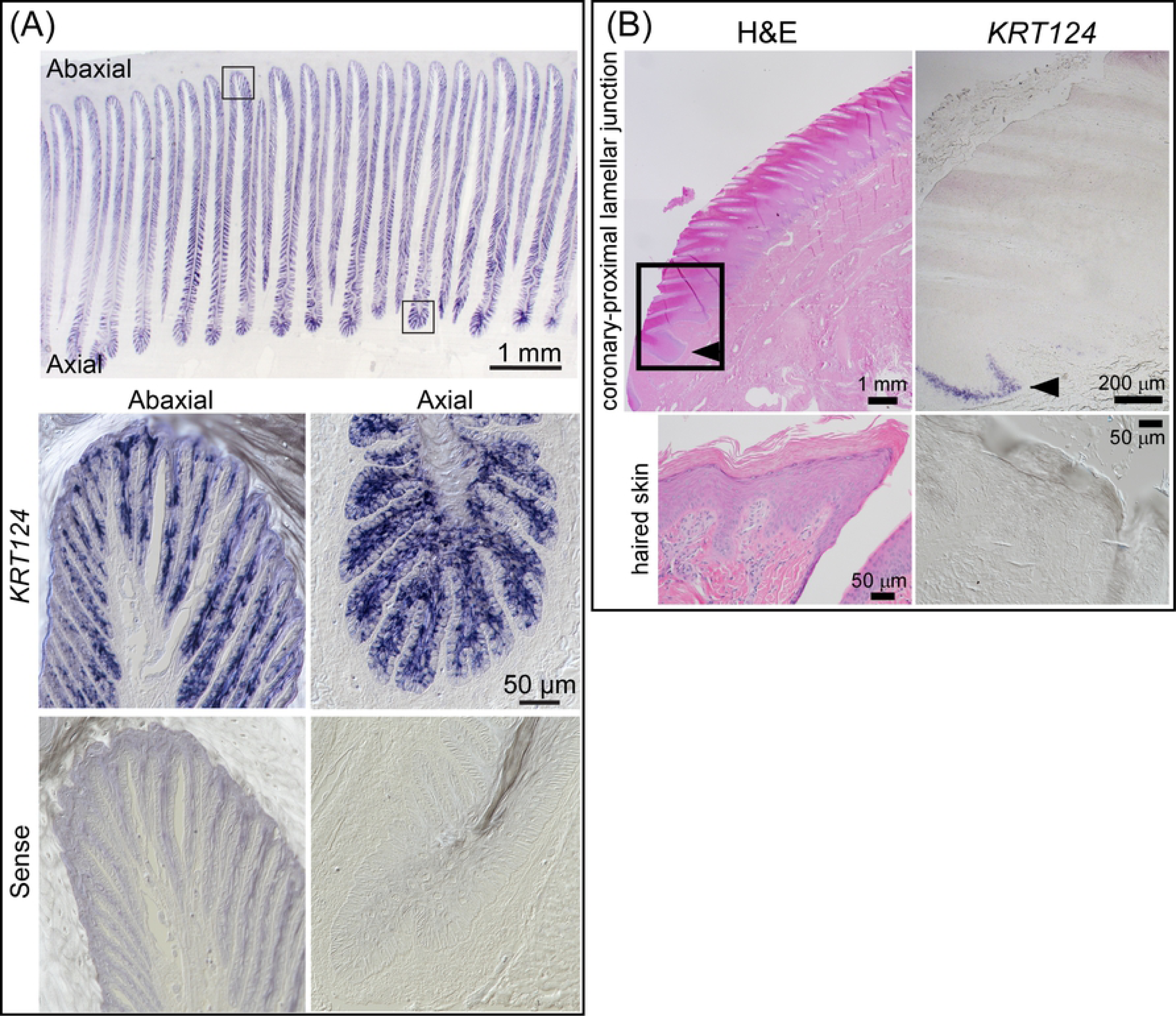
*KRT124* mRNA localizes to secondary epidermal lamellae and is absent from hoof coronet. **(A**) Representative images of *KRT124* localization to secondary epidermal lamellae (SELs) by in situ hybridization. *KRT124* localizes to suprabasal cells and, with less intense staining, to basal cells in all regions along the lamellae. Bottom panels: Representative differential interference contrast images of *KRT124* sense probe in situ hybridization shown as negative control. Axial (left) and abaxial (right) lamellar regions shown, corresponding to the axial and abaxial regions shown in top panel. Scale bar (50 μm) applies to all four lower panels. (**B**) Representative H&E and *KRT124* in situ hybridization images of a longitudinal section of the coronet and proximal lamella (arrowhead) and haired skin. Area of coronary-lamellar junction similar to the boxed area in the H&E image shows the abrupt transition from *KRT124*-negative keratinocytes in the coronary epithelium to *KRT124*-positive keratinocytes in a proximal lamella. All studies: n=3 using samples from 3 horses.

Although *KRT124* ISH staining is apparent in both basal and suprabasal cells in some regions (Fig 3A), it was generally more intense in the suprabasal cells and lighter or absent from basal cells (Fig 4). *KRT14* ISH, in contrast, is restricted to basal cells in lamellar tissue (Fig 4), similar to the previously reported localization of K14 protein in healthy lamellar tissue [16;19;24].

**Fig 4:**
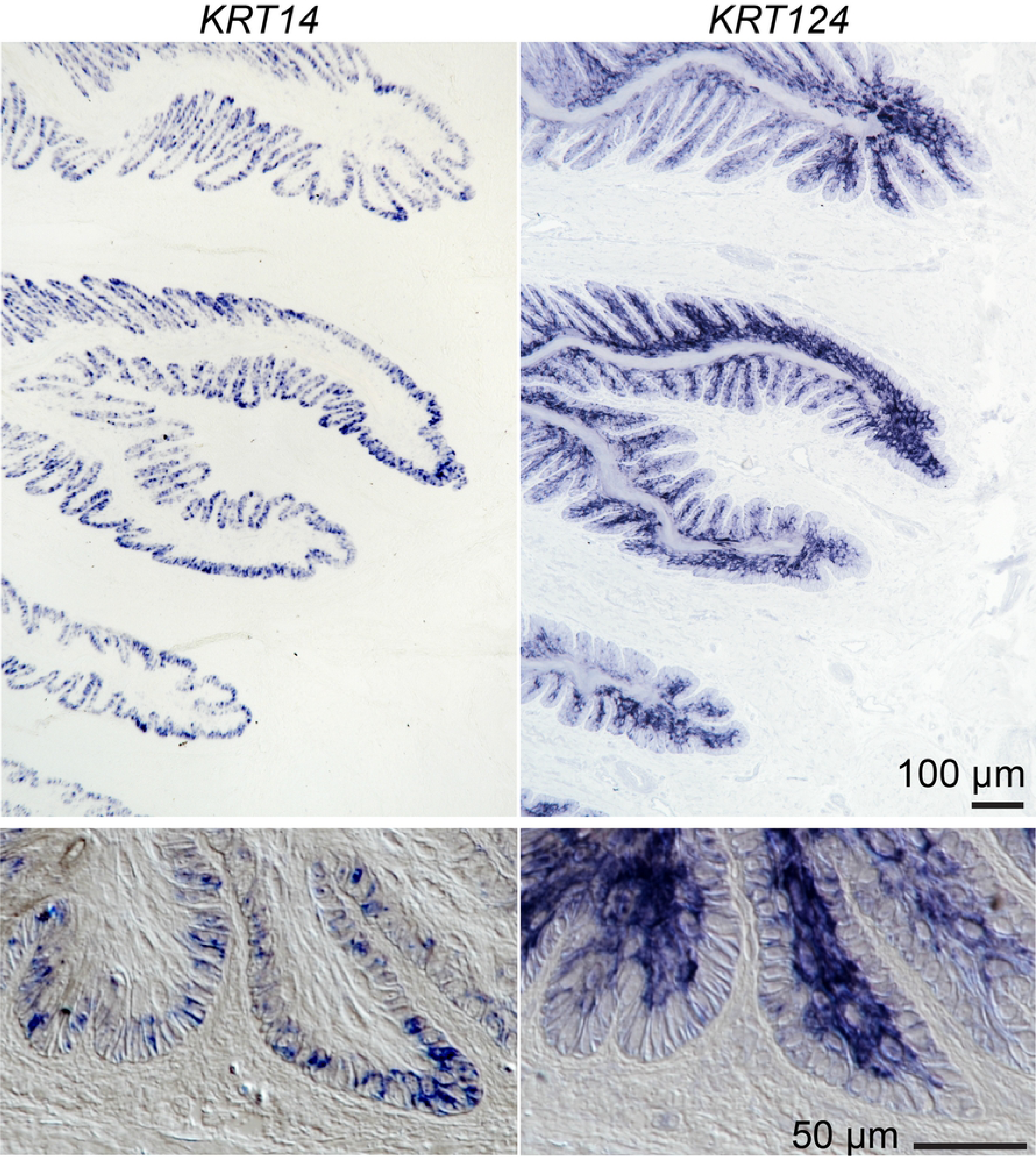
*KRT42* and *KRT124* mRNA localizes to secondary epidermal lamellae and is absent from hoof coronet. Representative images from in situ hybridization of *KRT14* (left) and *KRT124* (right) expression in serial sections. *KRT14* is restricted to basal cells, while *KRT124* is expressed primarily in suprabasal layers. Boxed regions marked on low magnification images are shown below at higher magnification and include differential interference contrast optics. All studies: n=3 using samples from 3 horses.

### 3.3 K42 and K124 monoclonal antibodies specifically detect hoof lamellar proteins on immunoblots

Monoclonal antibodies were generated against full-length recombinant equine K42 and against two different peptides from K124, as described in the Materials and Methods section, and characterized by immunoblotting (Fig 5). As shown in Fig 5A, the anti-K42 mAb detects a single band from lamellar tissue extract at the expected relative molecular mass (50 kDa), and comigrates with the second most abundant type 1 keratin in lamellar tissue, K14 [18]. Three anti-K124 mAb clones, K124A (against an N-terminal peptide), K124C, and K124D (the latter two against a single C-terminal peptide) were immunoreactive for a single major band at the expected relative molecular mass (54 kDa) in lamellar tissue extract (Fig 5A). The K14/K42, and K124 immunoblot bands correspond to two major protein bands that are visible by protein stain, even at the low total protein loads used for these studies (Fig 5A), as previously reported [18]. Anti-K124 C-terminal peptide clones K124C and K124D also detected a lower relative molecular mass minor doublet band that is not visible by protein staining in some lamellar tissue samples under these immunoblotting conditions (Fig 5B).

**Fig 5:**
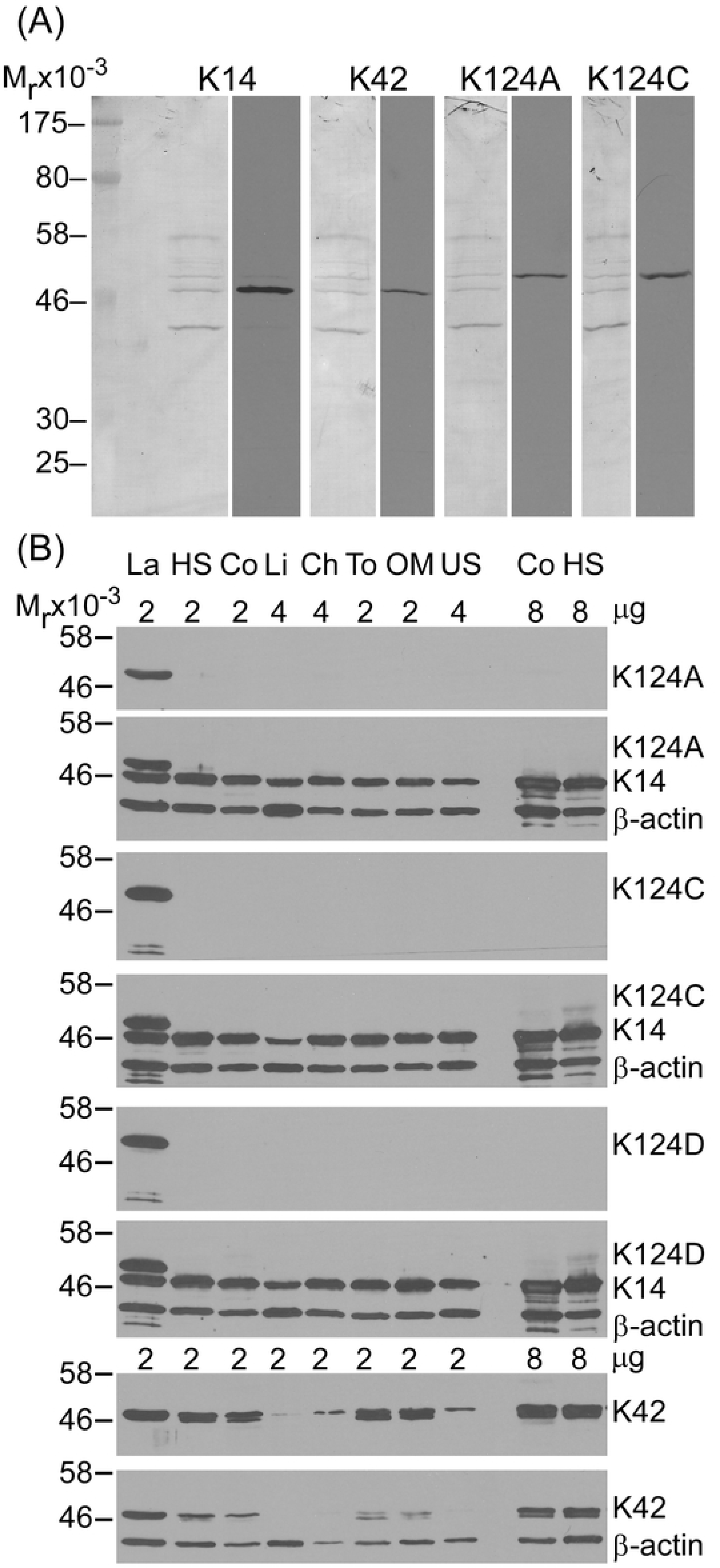
Detection of K14, K42, and K124 by immunoblotting with monoclonal antibodies. Representative immunoblots using mouse monoclonal anti-K14, anti-K42, or anti-K124, clones A, C, or D followed by secondary goat-anti-mouse-horse radish peroxidase (HRP) (n=5 using samples from 3 horses). (**A**) Representative images of K14, K42, K124A, and K124C immunoblot strips (right images) and amido black stain for protein of each blot (left images) from the same SDS-PAGE gel with 2 μg lamellar protein loaded per lane. K14, K42, K124A and K124C immunoblots detect a single band at the expected relative molecular weight in lamellar tissue. K14 and K42 co-localize to 50 kDa band and the K124 mAbs are immunoreactive with a 54 kDa band. (**B**) K14, K42, and K124 immunoblots of epidermal and surface epithelial tissue extracts demonstrate the specificity of the K124 mAbs to lamellar tissue. La: lamellar; HS: haired skin; Co: coronary; Li: Corneal limbus; Ch: chestnut; To: tongue; OM: oral mucosa; US: unhaired (glabrous) skin. Total protein load per lane indicated above tissue labels. K124 and K42 immunoblots reprobed with K14 and β-actin (K124 blots) or β-actin alone (K42 blots) without stripping to demonstrate equal load. K42 mAb detects a single band in lamellar, chestnut, and unhaired skin tissues and a doublet band in haired skin, coronary, tongue, and oral mucosa tissues. All three K124 mAbs detect a single major band and, for K124C and K124D, an additional, lower relative molecular mass minor doublet band only in lamellar tissue. Increased protein load confirms negative K124 mAb cross-reactivity to keratins in coronet and haired skin (last two lanes).

Immunoblotting with multiple stratified epithelial tissues was performed to evaluate antibody cross-reactivity (Fig 5B). The anti-K42 mAb detects a single band in lamellar, chestnut, and unhaired skin tissues and a doublet band in haired skin, coronary, tongue, and oral mucosa tissues. Immunoreactivity to non-lamellar tissues is less apparent than immunoreactivity to lamellar tissue following additional washes, but still clearly present for haired skin, coronary, tongue, and oral mucosa, consistent with antibody cross-reactivity at this antibody concentration (lower blot, Fig 5B). All three anti-K124 mAbs show no cross-reactivity to any of the non-lamellar stratified epithelial tissues tested. Increased protein load confirmed negative anti-K124 mAb cross-reactivity to keratins in coronet and haired skin (last two lanes for each immunoblot). Immunoblots with affinity-purified anti-K124, clones A and C, detected a single 54 kDa band with an antibody dilution of 1:5,000 and as little as 25 ng total lamellar protein load (data not shown).

### 3.4 K124 monoclonal antibodies specifically localize to hoof epidermal lamellae

As shown in Fig 6, indirect immunofluorescence using the anti-K124C mAb on cryosections demonstrates localization of K124 to the epidermal lamellae in a pattern that resembles that obtained by *KRT124* ISH (Fig 3-4). K124 localizes to suprabasal cells, and to a lesser degree, basal cells of all secondary epidermal lamellae. The keratinized axes of the primary epidermal lamellae are negative. Anti-K124C did not show any specific immunoreactivity to coronet or haired skin. Negative control parallel-run experiments on serial lamellar tissue cryosections showed some non-specific staining or autofluorescence of red blood cells, but no specific anti-K124C immunoreactivity (S1 Fig). The anti-K124A mAb showed some cross-reactivity to coronet and haired skin by indirect immunofluorescence and was not fully characterized for indirect immunofluorescence (data not shown). Preliminary indirect immunofluorescence studies with anti-K124D showed result similar to those with anti-K124C (data not shown).

**Fig 6:**
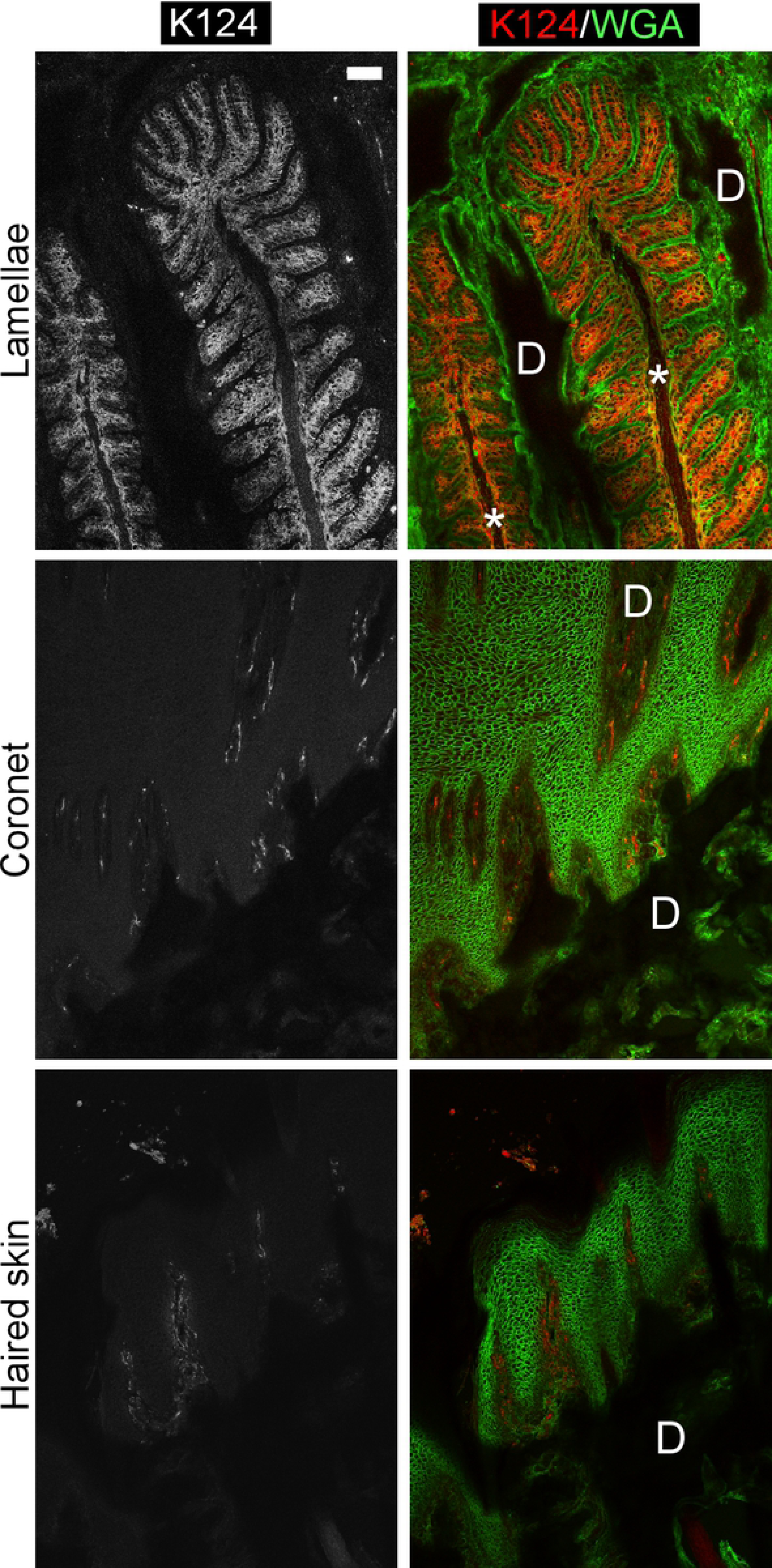
Localization of K124 to basal and suprabasal secondary epidermal lamellar cells by indirect immunofluorescence. Paraformaldehyde-fixed/sucrose-dehydrated/OCT-embedded cryosections from lamellar, coronet, and haired skin frozen tissues subjected to indirect immunofluorescence using the K124C mAb and fluorescein-conjugated wheat germ agglutinin (WGA) as a counterstain (n=3 using samples from 3 horses). K124C localizes to suprabasal cells, and to a lesser degree, basal cells of all SELs. Coronet and haired skin show negative staining for K124, with some autofluorescence of red blood cells visible in dermal tissues. (D): dermis; *: Keratinized axis of PEL. Scale bar = 20 μm. The same image adjustment to enhance red channel for ease of viewing was applied to all images.

## 4. Discussion

To our knowledge, this is the first report of the use of isoform-specific antibodies to localize a nail unit-specific keratin isoform (K124) in any species. In addition, this is the first demonstration that the expression of K124 is restricted to the equine (*E. caballus*) hoof lamellae, the highly folded inner epithelium of the hoof capsule, which is homologous to the nail bed of primates, rodents, and other species, and is absent from the germinative (“coronary”) region of the proximal hoof wall, which is homologous to the nail matrix [21;35;36]. K124 is the most abundant type II keratin of lamellar tissue [18] and, shown here (Fig 3, 4, 6), its expression is increased in suprabasal, compared to basal, lamellar keratinocytes, suggesting it is a terminal differentiation marker for these cells.

*KRT124* and *KRT42* were recently identified by total RNA sequencing from canine and equine skin, suggesting that transcripts for these keratins are found in equine haired skin, although this result was not validated by any complementary methods [20]. We amplified neither *KRT124* nor *KRT42* from haired skin total RNA by RT-PCR (Fig 2), nor did we localize K124 to haired skin by ISH, immunoblotting, or indirect immunofluorescence histology (Fig 3-6), using samples from multiple horses of several breeds (S1 Table). It is possible that the discrepancy relates to the anatomic location of the skin samples used since we collected samples from the dorsal region of the digit and the location of the skin biopsy used by Balmer, et al., is not specified [20]. Further investigation of equine skin keratin isoform expression is required to resolve this issue.

The rodent ortholog of *KRT124, Krt90* (formerly *Kb15*) and the opossum ortholog (*Kb15*) have not been characterized beyond genomic mapping and identification as likely functional genes in those species [22;23]. However, based on the relative protein amounts of K42 and K124 in lamellar tissue, we had previously suggested that these keratins hybridize in lamellar tissue, and are therefore expected to co-localize in the hoof lamellae [18]. Murine *Krt42* (formerly *K17n*) expression has been localized to the nail matrix and nail bed and functional canine *KRT42* and *KRT124* genes have been identified, suggesting that the nail bed expression of K42 and K124/K90 may be conserved across mammalian and marsupial species, but was lost from the thinner and non-weight-bearing nail units of primates, where both keratins exist only as pseudogenes [20-22]. Equine *KRT42* and *KRT124* have a more restricted tissue localization than murine *Krt42* since the latter is expressed in the nail matrix [22], but the former are not expressed in the homologous coronary region of the hoof wall, as assessed by RT-PCR for *KRT42* (Fig 2) and multiple methods for *KRT124*/K124 (Fig 2-6). The isoform-specific anti-K124 mAbs described here may allow protein localization and tissue distribution of K124/K90 in other species.

The biology of equine lamellae is also of interest due to the prevalence of equine laminitis, a common and devastating disease affecting this tissue. Laminitis results in epidermal pathologies that include abnormal hyperplastic and acanthotic epidermal tissue [37], epidermal dysplasia and metaplasia, loss of cell adhesion, apoptosis, and necrosis[27], and expression of cellular stress, activation, and altered differentiation markers [24;38-40]. Similar nail abnormalities involving the nail bed were recently described in association with ageing in several inbred strains of mice [41]. Our anti-K124 mAbs will be useful for the investigation of histopathological changes in lamellar and nail bed keratin expression and as a tissue-specific differentiation marker for *in vitro* studies. Keratins, as the most abundant proteins and as epithelial-specific proteins, are useful biomarkers of epithelial cell stress, apoptosis and necrosis in several human diseases, including various carcinomas [42] and several types of liver disease [43]. K124 could similarly serve as a tissue-specific disease biomarker for equine laminitis and nail unit disease in other species that express it.

In conclusion, we have characterized the expression of keratin isoforms that specifically localize to the highly specialized inner epithelium of the equine hoof capsule. For the first time, we have generated and characterized nail unit-specific anti-K124 mAbs, which localize specifically to the secondary epidermal lamellae and do not cross-react with proteins from several stratified epithelial tissues. We suggest that these hoof-specific keratins are essential components of the equine suspensory apparatus of the distal phalanx and provide the mechanical properties of strength and elasticity that enable single digit, unguligrade locomotion in the equidae, a signature evolutionary adaptation of this genus.

## 5. Acknowledgments

CDSS and LC are grateful to Micheal Layden, Jamie Havrilak and Dylan Faltine-Gonzalez for sharing their expertise, equipment and reagents for in situ hybridizations. HGH thanks Julie Engiles, Sue Lindborg, Renata Linardi, Mark Modelski and Susan Megee for their work on the tissue bank.

## 7. Supporting Information

**S1 Table: Breed, age, and sex of horses (*E. caballus*) used in experiments**

**S1 Fig: Indirect immunofluorescence negative control for lamellar tissue.** Lamellar tissue cryosection, serial to the one shown in Fig 6, subjected to indirect immunofluorescence and fluorescein-conjugated wheat germ agglutinin (WGA) as a counterstain, omitting the K124C mAb to show non-specific staining (n=3 using samples from 3 horses, representative image shown). (A) Red channel, secondary antibody alone (white). (B) Secondary antibody alone (red) and fluorescein-WGA counterstain (green). Scale bar = 20 μm. The same image adjustment to enhance red channel for ease of viewing as that applied to Fig 6 was applied to these images.

